# Modulation of circadian rhythms in articular cartilage by heat pulses

**DOI:** 10.1101/2024.01.26.577368

**Authors:** Cátia F. Gonçalves, Michal Dudek, Dharshika RJ Pathiranage, Anna Paszek, Zeyad El-Houni, Shiyang Li, Gwenllian Tawy, Leela Biant, Judith A. Hoyland, Qing-Jun Meng

## Abstract

**Objective:** Prior studies have shown that disruption of the circadian clock leads to cartilage degeneration in mice while shift work is associated with higher risk of osteoarthritis (OA) in humans. In this study we investigated the potential of heat pulses to restore dampened circadian rhythms in articular cartilage.

**Methods:** Femoral head cartilage explants and primary chondrocytes were isolated from PER2::LUC mice. Human femoral condyle cartilage was obtained from osteoarthritic patients undergoing total knee replacement. Tissues and cells were exposed to heat shock at various temperatures (37-43 °C) and incubation lengths. Bioluminescence from explants and cells was recorded in real-time. RNA sequencing and qPCR were used to assess gene expression changes in response to heat.

**Results:** We established that a 60-min pulse at 43 °C was sufficient to restore dampened PER2::LUC rhythms in mouse cartilage explants or in primary chondrocytes. Transcriptome analysis in mouse articular cartilage showed an up-regulation of genes encoding heat shock proteins and collagens, and a transient down-regulation of *Sox9*, *Runx2*, *Per1*, *Clock* and *Cry2*. Heat induced the expression of circadian clock genes in human osteoarthritic knee cartilage. Mechanistically, inhibition of HSP90 activity or perturbation of F-actin polymerisation blocked the heat-induced resynchronisation of circadian rhythms.

**Conclusion:** Together, these data have contributed to a greater understanding of the multifaceted nature of the connections between circadian timekeeping, heat stress responses and homeostasis in articular cartilage. These findings also suggest that time-prescribed temperature increases could be developed into a non-invasive intervention to slow down tissue ageing and restore homeostasis in osteoarthritic joints by improving circadian oscillations of cartilage rhythmic pathways.

## Introduction

Environmental temperature cycles are a primordial resetting cue for circadian rhythms in all living systems, with the exception of homeothermic vertebrates [1]. Although mammals do not usually entrain to environmental temperature cycles [2], the circadian rhythms of body temperature, driven by the suprachiasmatic nucleus (SCN), function as a universal time cue for the entrainment of autonomous circadian oscillators throughout the body [3]. On the molecular level the circadian clock mechanism is composed of transcription factors CLOCK and BMAL1 and their repressors PER and CRY forming a negative feedback loop that generates 24-hour rhythms in the expression of genes and proteins.

Many aspects of cartilage physiology, such as the pathways involved in its maintenance and repair, are controlled by the circadian clock [4–6]. Disruption of circadian rhythms has been associated with pathophysiological changes in cartilage in both mice and humans [6,7]. Moreover, in ageing mice circadian rhythms in cartilage exhibit reduced amplitude [4].

Interestingly, circadian rhythms in xiphoid cartilage and intervertebral disc can be entrained by cyclic temperature changes that approximate daily body temperature fluctuations (38.5 °C for 12 hours/36.0°C for 12 hours) [4,6]. Other studies have also demonstrated that short-term, high-amplitude temperature pulses can entrain circadian rhythms in fibroblasts [8,9], and enhance oscillation amplitude in liver and lung explants [10]. This raised the hypothesis that temperature pulses can be utilised to ameliorate dampened circadian rhythms in cartilage and improve the clock-controlled regulation of homeostatic processes in the tissue, which could have potential implications in the context of diseases such as osteoarthritis.

Importantly, the mechanistic understanding of how temperature cycles entrain circadian clocks is still limited. For example, it is known that the heat shock response is involved in the regulation of the core circadian clock, affecting the circadian phase and period in multiple mouse tissues [3,11,12]. Data have also supported that Heat shock factor 1 (HSF1) induces the transcription of *Per2* after heat shock and that the stability of BMAL1 is affected by HSP90 [8,13]. Heat stress can also lead to the reorganisation and even collapse of actin networks [14,15]. Actin is an essential component of the cytoskeleton that participates in the regulation of cell shape, movement, division, and in cell signalling [16]. Particularly, sensing of the extracellular matrix stiffness through actin cytoskeleton with involvement of Rho/ROCK (Rho-associated protein kinase) and SRF (Serum response factor) pathways has been shown to affect the circadian clock [17–19].

In this study, the optimal temperature and duration to restore dampened PER2::LUC rhythms were established in mouse articular cartilage from both young and ageing mice, and in primary chondrocytes. The changes in the expression of circadian clock genes were validated in human osteoarthritic cartilage explants. Transcriptome analysis and pharmacological approach were used to reveal the downstream pathways and underlying molecular mechanisms for the heat-induced resynchronisation of circadian rhythms.

## Methods

### Animal husbandry

All experimental procedures were carried out in accordance with the Animals (Scientific Procedures) Act of 1986. Mice were group housed under a 12 hours light/12 hours dark regimen (lights on at 7 AM and off at 7 PM) with *ad libitum* access to standard rodent chow and water. All animals were bred by the Biological Services Facility at the University of Manchester and euthanised by a schedule 1 procedure between *zeitgeber* time 5.5 (ZT5.5) and ZT6.5 (between 12.30 PM and 1.30 PM). The time at which lights are switched on is herein described as ZT0 and lights off as ZT12. PER2::LUC mice carrying the firefly luciferase gene fused in frame with the 3’ end of *Per2* were utilised in all studies [20].

### Articular cartilage explants

Articular cartilage from one- to 12-month-old mice was obtained by careful dissection of the femoral head articular cartilage [6]. Explants were maintained in Dulbecco’s Modified Eagle Medium/Nutrient Mixture F-12 (DMEM/F-12, Gibco) supplemented with 10 % foetal bovine serum (FBS, Gibco) and 100 units penicillin/100 μg/mL streptomycin (Sigma-Aldrich).

### Primary articular chondrocytes

Primary articular chondrocytes were isolated from five-day-old mice as previously described [21] with minor modifications. In brief, articular cartilage tissues were isolated from femoral heads, femoral condyles and tibial plateaus of mice from one litter and pooled. Explants were rinsed with 1 × phosphate buffered saline (PBS, Sigma-Aldrich) and incubated in digestion solution (3 mg/mL collagenase D [Roche] in DMEM/F-12 [Gibco]) for 45 min at 37 °C with intermittent vortex mixing to dislodge soft tissues. Subsequently, samples were diced using a sterile scalpel and incubated overnight at 37 °C in the digestion solution. Cell aggregates were dispersed by successively passing the solution through 25-, 10- and 5-ml Pasteur pipettes. The cell suspension was filtered through a 70 μm cell strainer (EASYstrainer, Greiner Bio-One) and then centrifuged for 10 min at 400 ×g. Finally, the pellet was resuspended in DMEM/F-12 (Gibco) supplemented with 10% FBS (Gibco), and 100 units penicillin/100 μg/mL streptomycin (Sigma-Aldrich). Primary articular chondrocytes were used up to passage 2.

### Bioluminescence experiments

#### Culture conditions

For bioluminescence recording, articular cartilage explants or primary articular chondrocytes were cultured in DMEM/F12 D2906 media (Sigma-Aldrich) supplemented with 10 % FBS (Gibco), 1.2 g/L sodium bicarbonate (Sigma-Aldrich), 100 units penicillin/100 μg/mL streptomycin (Sigma-Aldrich) and 100 μM luciferin (Promega). Chondrocytes were incubated in the presence of 100 nM of dexamethasone for 60-min prior to the media change. Samples were maintained in 35 mm cell culture dishes (Corning) sealed with 40 mm glass cover slips (Thermo Fisher Scientific) and high vacuum grease (Dow Corning). Sealed dishes were placed in a LumiCycle luminometer (ActiMetrics) and PER2::LUC bioluminescence was recorded continuously.

#### Temperature pulses

LumiCycle apparatuses were housed inside cell culture incubators maintained at 37 °C. For temperature pulses, sealed dishes were removed from the LumiCycle and placed in cell culture incubators at 40 °C or 43 °C for 30 min, 60 min or 90 min; no-treatment and dexamethasone-treated (100 nM, Sigma-Aldrich) controls were moved to an incubator at 37 °C for the duration of the treatment. After treatment, all dishes were returned to the luminometer set at a constant temperature of 37 °C. Temperature pulses were performed either at a projected PER2::LUC peak or trough on the third, fourth or fifth day after bioluminescence recordings were initiated. The time of the projected peak or trough was calculated by adding the period in hours to the time of the second PER2::LUC peak or trough.

#### Treatment with IL-1α

PER2::LUC rhythms were monitored for 42 hours prior to the addition of 10 ng/mL of IL-1α (R&D) or matching vehicle (UltraPure DNase/RNase-Free Distilled Water, Invitrogen). Explants were heat shocked immediately or three hours after IL-1α addition.

#### Data analysis

LumiCycle Analysis software version 2.60 (ActiMetrics) was used to analyse bioluminescence data. Raw data was detrended by subtracting background bioluminescence using a 24-hour moving average. The period was determined by fitting this baseline-subtracted data to a damped sine wave [“LM Fit (Damped sin)]. Phase shifts were measured by subtracting the time of observed peaks in treatment samples to the time of observed peaks in control samples.

#### Statistical analysis

GraphPad Prism version 9.3.1 (GraphPad Software, Inc.) was used for all statistical analysis. Assumption of normality was confirmed using the Shapiro-Wilk test (α = 0.05) in all analysed datasets. The statistical tests performed are reported in figure legends.

### RNA-sequencing

#### Experimental design

Articular cartilage from one-month-old mice was obtained by careful dissection of the femoral head articular cartilage [6]. Explants were cultured in DMEM/F12 media (Sigma-Aldrich) supplemented with 10 % FBS (Gibco), 1.2 g/L sodium bicarbonate (Sigma-Aldrich), 100 units penicillin, 100 μg/mL streptomycin (Sigma-Aldrich) and 100 μM luciferin (Promega). Samples were maintained in 35 mm dishes sealed with 40 mm glass cover slips (Thermo Fisher Scientific) and high vacuum grease (Dow Corning). Sealed cell culture dishes were placed in an incubator at 37 °C for approximately five days. Samples were collected at four time points: before heat shock at a projected trough or peak (Time 0), immediately after the 60-min pulse at 43 °C (Time 1), and one and three hours after the pulse (Time 2 and Time 4, respectively).

#### RNA extraction from mouse articular cartilage

Frozen tissue was reduced to a powder in the presence of 500 μL of TRIzol (Invitrogen) in a liquid nitrogen-cooled Braun Mikro-Dismembrator Vessel (B. Braun Biotech International). Another 500 μL of TRIzol was added to the powdered tissue and incubated at room temperature for 5 min to allow for complete dissociation of nucleoprotein complexes. 200 μL of chloroform (Fisher Chemical) was added to this suspension, mixed vigorously, and centrifuged at 12,000 ×g for 15 min at 4 °C. 350 μL of the upper aqueous phase was transferred to a new tube, mixed vigorously with 400 μL TRIzol and 200 μL chloroform, and centrifuged at 12,000 ×g for 15 min at 4 °C. Then, 450 μL of the upper aqueous phase was transferred to a new tube, mixed vigorously with 200 μL TRIzol and 100 μL chloroform, and centrifuged at 12,000 ×g for 15 min at 4 °C. Lastly, 185 μL 3.6 M sodium chloride (Fisher Chemical)/2.4 M sodium acetate (Millipore Sigma-Aldrich), 10 μg RNA grade glycogen (Thermo Fisher Scientific) and 700 μL isopropanol (Fisher Chemical) were added to 500 μL of the upper aqueous phase, mixed vigorously, incubated at room temperature for 15 min, and centrifuged at 12,000 ×g for 30 min at 4°C. Supernatant was discarded, pellet was washed three times with 500 μL of 75 % ethanol (Fisher Chemical), and centrifuged for at 12,000 ×g for 5 min at room temperature. The RNA pellet was dried on a heat block for 2 min at 50 °C. 10 μL of RNAse-free water (Sigma-Aldrich) were added to dissolve the pellet; the yield was quantified using the Qubit fluorometer (Invitrogen) and the absence of genomic DNA contamination was checked using the TapeStation automated electrophoresis system (Agilent). RNA samples were stored at -80 °C until further processing.

#### Library construction and sequencing

mRNA library preparation and sequencing were prepared at the University of Manchester’s Genomic Technologies Core Facility. Libraries were generated using the TruSeq Stranded mRNA assay (Illumina) according to the manufacturer’s protocol. Libraries were sequenced on the NextSeq500 instrument (Illumina) using the NextSeq 300-cycle high output kit (Illumina) to generate 2 × 151 bp paired-end reads.

### Bioinformatics analysis

#### Read pre-processing, filtering and mapping

Quality control and pre-processing of FASTQ files were performed using fastp v0.20.0 [22] with the polyA trimming and cut-right settings enabled. Reads were then mapped to the Ensembl GRCm38 mouse genome and GENCODE M23 annotation using STAR v2.5.3a with ENCODE settings and the index generated by RSEM v1.3.0 [23,24].

#### Differential gene expression analysis

Quantification of transcripts was performed using RSEM v1.3.0 [24]; and differentially expressed genes were identified with the DESeq2 v1.28.1 package from Bioconductor in R [25]. To account for possible sources of variation a “condition” term (P0, P1, P2, P4, T0, T1, T2 and T4) and a “biological replicate” term (A, B and C) were fitted with a generalised linear model. The full model was compared to a reduced model (∼condition+replicate). Differentially expressed genes (DEGs) were tested based on their fold change and adjusted *p* value corrected for multiple comparisons via the Benjamini-Hochberg method [26]. The least binary logarithmic fold change threshold was set to more than one (lfcThreshold, θ > 1) and adjusted *p* value was set to a false discovery set of less than 0.05 (alpha, α < 0.05).

Principal components analysis was performed with the DESeq2 v1.28.1 package on variance stabilised counts [25]. Because the variance stabilising transformation does not use the design model to remove variation in the data, the removeBatchEffect function from limma v3.44.1 was employed to remove any shifts in the log2-scale expression data that could be explained by batch effects [27].

To allow the visualisation of expression changes at the gene level, the standardize function from the easystats v0.2.0 package was employed to obtain Z-scores from the batch-corrected variance-stabilised counts. Next, unsupervised hierarchical clustering was performed using the pheatmap v1.0.12 package with dissimilarity metrics based on Pearson’s correlation, complete linkage as the agglomeration method and k-means clusters [28]. The number of clusters was defined by visual inspection of the heatmap; this was complemented by plotting the clustered trajectories of gene expression and the k-means centroids for each cluster with ggplot2 v3.3.3. To identify over-represented genes, GO terms associated with the different clusters the WEB-based GEne SeT AnaLysis Toolkit (WebGestalt) was utilised; only terms with a FDR less than 0.05 were selected and redundancy reduction was accomplished via the affinity propagation algorithm [29].

### Human osteoarthritic knee cartilage

Human knee articular cartilage samples were obtained under National Health Service Research Ethics Committee approval (REC Reference: 22/LO/0102) with prior written informed consent. Patients were eligible if they were listed for a total knee replacement to treat end-stage osteoarthritis. All patients provided informed consent for the study prior to their surgery. At removal intra-operatively, tissue explants from the femoral condyles were cultured in DMEM high glucose (D6429, Sigma) supplemented with 10 % foetal bovine serum (Sigma), 100 units penicillin/100 μg/mL streptomycin (Sigma) and 40 units nystatin (Sigma).

| Donor | Age (years) | Sex | Knee |
| --- | --- | --- | --- |
| A | 65 | Male | Right |
| B | 72 | Female | Left |
| C | 67 | Male | Left |

### Reverse transcription and quantitative polymerase chain reaction (RT-qPCR)

mRNA from cartilage samples was extracted using the same method described above in mouse RNA sequencing. Complementary DNA (cDNA) was generated from 400 ng of total RNA using the high-capacity RNA-to-cDNA kit (Applied Biosystems) according to the manufacturer’s instructions. RT-qPCR was performed in a total volume of 10 μL, comprising 5 μL of TakyonTM MasterMix (Eurogentec), 0.5 μL of TaqMan® gene expression assay probes (Table 1), 1 μL cDNA template at a concentration of 10 ng μL-1 and 3.5 μL of nuclease-free water (Sigma-Aldrich). No template controls were included for all reactions. RT-qPCR was carried out on a StepOne Real-Time PCR System (Applied Biosystems) according to the following programme: 2 min at 50 °C to prevent carryover, 3 min at 95 °C to activate the TakyonTM enzyme, and 40 cycles of denaturation at 95 °C for 30 sec and annealing/extension at 60 °C for 60 sec. Gene expression data was obtained using the ΔΔCt method in which samples were first normalised to the mean levels of the reference genes *CANX* and *CSDE1* in the case of human samples [30] and *Rpl13a* and *Ywhaz* in the case of mouse samples [31]. Relative normalisation was performed in relation to the time-matched NTC control in human samples and to time 0 in mouse samples.

List of Taqman gene expression assay probes.

List of Taqman gene expression assay probes.
| Gene | Assay identity |
| --- | --- |
| *BMAL1* | Hs00154147_m1 |
| *CANX* | Hs01558409_m1 |
| *CLOCK* | Hs00231857_m1 |
| *COL2A1* | Hs00264051_m1 |
| *COL2A9* | Hs00895570_m1 |
| *CRY1* | Hs00172734_m1 |
| *CRY2* | Hs00323654_m1 |
| *CSDE1* | Hs00918650_m1 |
| *CXXC5* | Hs00212840_m1 |
| *HSPA1A* | Hs00359163_s1 |
| *HSPA1B* | Hs01040501_sH |
| *HSPB1* | Hs00356629_g1 |
| *JARID2* | Hs01004457_m1 |
| *LNPEP* | Hs00893646_m1 |
| *PER1* | Hs01092603_m1 |
| *PER2* | Hs00256143_m1 |

## Results

### Heat pulses restore dampened circadian rhythms in mouse articular cartilage and chondrocytes

Femoral head cartilage from PER2::LUC reporter mice were monitored in *ex vivo* culture by real-time bioluminescence recording for approximately five days, at which point whole-tissue circadian rhythms are dampened due to desynchronisation. Next, heat pulses at 40 °C or 43 °C for 30, 60 or 90 min were applied either at a projected peak or trough of PER2::LUC. Control explants were incubated at 37 °C for the treatment duration either in the presence (positive control) or absence (negative control) of 100 nM dexamethasone, a glucocorticoid hormone analogue that induces circadian gene expression [32]. Incubation of cartilage explants at 43 °C for 60 min at projected peak of PER2::LUC resulted in the most effective increase in oscillation amplitude (Fig. 1A). A 90-minute exposure at 43 °C produced a comparable reaction and did not provide any additional advantage compared to a 60-minute incubation. All other conditions did not have any significant effect on the circadian rhythm (Supplementary Fig. 1A-D). Similar results were obtained with mouse primary chondrocytes: a 60-min pulse at 43 °C resulted in the highest amplitude increase. However, cells responded to the heat pulse similarly both at a projected peak or trough (Supplementary Fig 2 A-D). Repeating the heat pulses on explant tissues or primary chondrocytes every 24 hours for 3 days did not magnify the response beyond the amplitude of a single treatment; however, the explants did respond to each pulse with resynchronisation (Supplementary Figs. 3 A-E and 4 A-E).

**Figure 1.**
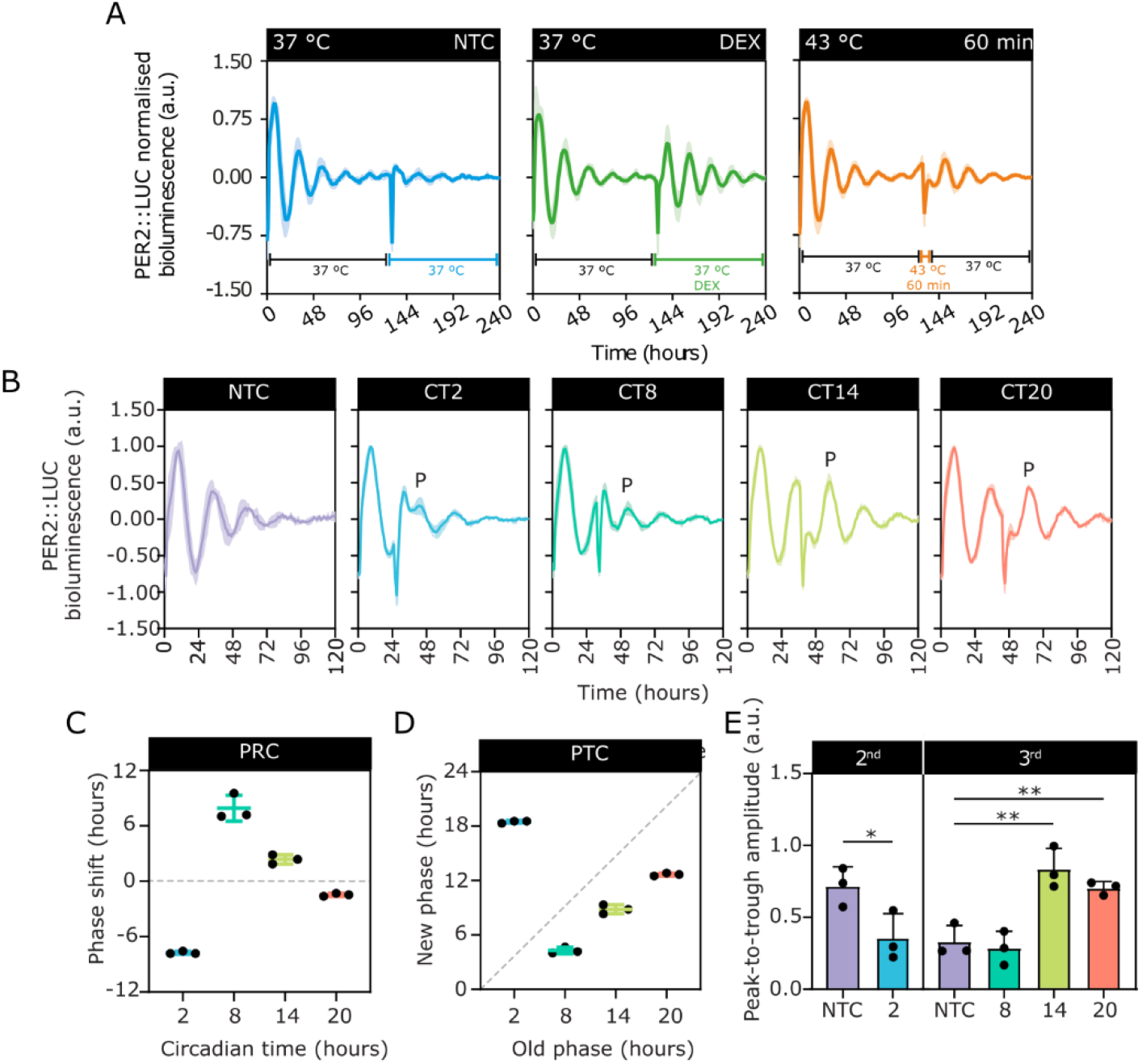
Dampened circadian rhythms in murine articular cartilage can be restored by heat pulses. Normalised PER2::LUC bioluminescence profiles of mouse femoral head articular cartilage. **A.** Explants were incubated for 60 min at 43 °C at a projected peak of PER2::LUC. NTC, no-treatment control; DEX, 100 nM dexamethasone. **B.** Tissues were incubated at 43 °C for 60 min at four time points after the first oscillation: circadian time 2 (CT2), CT8, CT14 and CT20. The first peak after treatment is highlighted by the letter P. **C**. Quantification of the phase response curve of articular cartilage in B. Phase shifts were determined by subtracting the time of the peak after treatment (P) to the time of observed peaks in NTC explants. **D**. Phase transition curve showing the phase of circadian rhythms in cartilage explants before (old phase) and after (new phase) the heat pulse. The dashed line segment represents a slope equal to one. **E**, Peak-to-trough amplitude of the first oscillation after heat shock (P) compared to the respective NTC oscillation as indicated. Second oscillation: two-tailed unpaired t-test [t(4) = 2.820, *p* = 0.0478]. Third oscillation: one-way ANOVA [F(3,8) = 18.06, *p* = 0.0006] and post-hoc Dunnett’s multiple comparison test (NTC vs. CT14, *p* = 0.0014; NTC vs.CT20, *p* = 0.0086). Data were reported as mean ± SD. n ≥ 3. CT2 data was compared to the second peak in the NTC, while CT8, CT14 and CT20 data was compared to the third peak.

To further elucidate the phase dependent effects of 60-min pulses at 43 °C, explants were heat shocked at four different circadian phases: circadian time 2 (CT2), CT8, CT14 and CT20 (Fig. 1B). The PER2::LUC peak occurred at CT9 and the trough at CT21. A heat pulse at CT2 produced a substantial phase delay (Mean = -7.737, SD = 0.144 hours), and at CT8 a phase advance of similar magnitude was observed (Mean = 7.910, SD = 1.397 hours). Less pronounced phase shifts (̴2 hours) were observed at CT14 and CT20 (Fig. 1B). The magnitude of the phase shifts observed at CT2 and CT8 as well as the slope of the phase transition curve close to 0 suggested a type 0 phase resetting (response to strong stimuli) of circadian oscillators in cartilage (Fig 1C-E). A similar type 0 resetting was also observed in primary chondrocytes with increased sampling frequency (Supplementary Figs. 4 F-H).

With increasing age, the cartilage circadian rhythm in mice shows reduced oscillation amplitude [4,33]. Therefore, we investigated whether the heat pulses can resynchronise cartilage tissue clock in ageing mice. As reported previously [4,34], cartilage explants from aged mice had lower starting amplitude compared to young mice. A 60-min pulse at 43 °C increased PER2::LUC oscillation amplitude in cartilage explants from both young adult and ageing mice to their respective starting amplitude (Fig. 2A, B), suggesting the circadian clock in ageing cartilage tissue remains responsive to heat stimulation (Fig. 2B).

**Figure 2.**
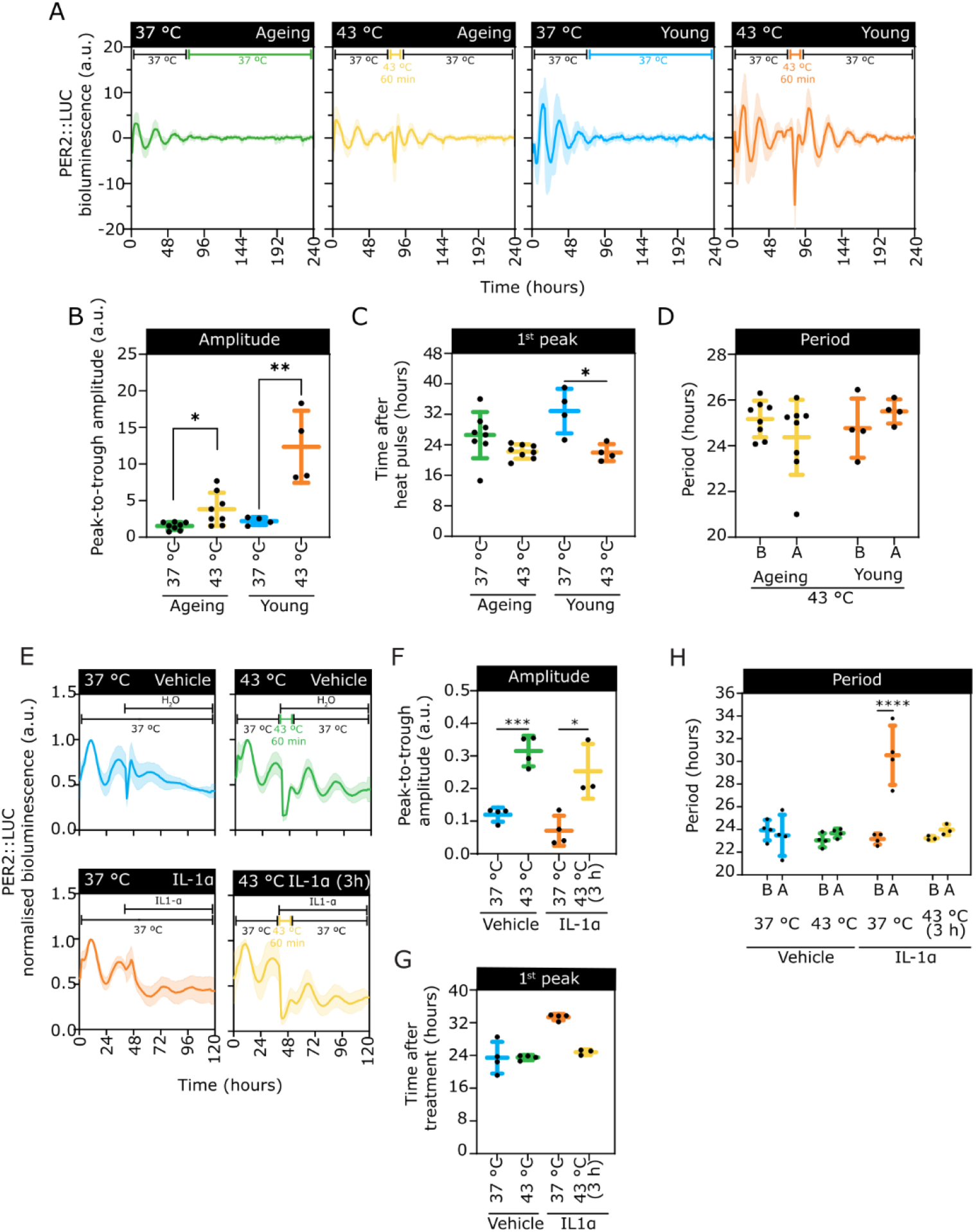
Heat pulse synchronises circadian clocks in ageing, and IL-1α treated mouse articular cartilages. **A.** Normalised PER2::LUC bioluminescence profiles of mouse femoral head articular cartilage. Tissues were exposed to the heat pulse at 43 °C for 60 min after 82 hours from the start of recording. **B**. Peak-to-trough amplitude of the first oscillation after the heat pulse compared to the respective control. Two-tailed unpaired t-test with Welch’s correction. Ageing: t(7.792) = 2.792, *p* = 0.0242; Young: t(3.073) = 4.095, *p* = 0.0252. **C**. Time of the first peak following heat treatment. Young: two-tailed unpaired t-test [t(6) = 3.488, *p* = 0.0130]. **D**. Period of the circadian rhythms before (B) and after (A) treatment. Data were reported as mean ± SD. n ≥ 4. **E.** Normalised PER2::LUC bioluminescence profiles of mouse femoral head articular cartilage. IL-1α (10 ng/mL) or vehicle control were added to recording media at 42 hours. Tissues were exposed to the heat pulse at 43 °C after a three-hour incubation. IL-1α remained in the recording media for the rest of the time course. **F**. Peak-to-trough amplitude of the first oscillation after the heat pulse compared to the control. Vehicle: two-tailed unpaired t-test [t(6) = 7.579, *p* = 0.0003]; IL-1α: one-way ANOVA [F (2, 7) = 7.862, *p* = 0.0162] and post-hoc Dunnett’s multiple comparison test (*, *p* = 0.0101). **G**. Time of the first peak following heat treatment. IL-1α: one-way ANOVA [F (2, 7) = 216.6, *p* < 0.0001] and post-hoc Dunnett’s multiple comparison test (****, *p* < 0.0001); IL-1α-37 °C vs. Vehicle-37 °C: two-tailed unpaired t-test with Welch’s correction [t(3.243) = 5.014, *p* = 0.0126]. **H**. Period of the circadian rhythms before (B) and after (A) treatment. Two-way ANOVA [F (1, 13) = 14.05, *p* = 0.0024] and Šídák’s multiple comparisons test (***, *p* < 0.0001). Data were reported as mean ± SD. n = 3.

In aged, injured or osteoarthritic joints, the expression of catabolic cytokines (e.g., TNF-α and IL-1) is elevated in both animal models and humans [35,36]. This aberrant expression of inflammatory mediators is known to elicit metabolic imbalances in chondrocytes in which catabolic responses overcome the attempt to repair [35,36]. IL-1 (and not TNF) has been shown to disrupt the circadian rhythm in cartilage and chondrocytes [37,38]; thus, we investigated whether heat pulses can counteract the effect of proinflammatory cytokines on the clock. PER2::LUC circadian rhythms were monitored for 42 hours prior to chronic treatment with 10 ng/mL of IL-1α (Figure 2C). The treatment resulted in dampened amplitude of oscillations and lengthening of the circadian period. Explants were heat shocked for 60 min at 43 °C three hours after IL-1α treatment. The heat pulse increased oscillation amplitude even in the presence of IL-1α, with no significant difference in circadian period before and after treatment (Fig. 2D), suggesting this approach could serve as an effective intervention to preserve circadian rhythms in the face of inflammation.

### The heat pulse elicited broad transcriptional changes in mouse articular cartilage

To gain more insight into the mechanism of clock synchronisation by the heat pulse and uncover potential beneficial homeostatic effects, we utilised RNA sequencing of mouse articular cartilage explants exposed to 43 °C for 60 min at multiple timepoints (before the heat pulse (Time 0), immediately after heat pulse (Time 1), and one (Time 2) and three (Time 4) hours after the pulse) (Fig. 3A). The heat pulse was applied either at a projected peak (when circadian clock responds with resynchronisation) or a projected trough of PER2::LUC (when resynchronisation does not occur). Principal Component Analysis (PCA) showed a clear segregation between timepoints after heat, but not whether the samples were treated at a projected peak or trough of PER2::LUC, indicating that time after the heat pulse is the biggest source of variance on a global scale in the dataset (Fig. 3B). Upon heat pulse, there was a predominant up-regulation of gene expression at both peak and trough treatments, with the number of up-regulated genes progressively increasing over time (Fig. 3C). In P1 vs. P0 there were approximately twice as many up-regulated genes than in T1 vs. T0 (Fig. 3C). Likewise, the number of down-regulated genes at P1 vs. P0 and P2 vs. P0 was far greater than at the trough (T1 vs. T0 and T2 vs. T0). At the peak, 809 statistically significant genes (FDR < 0.05) were shared between the three pairwise comparisons (Fig. 3D), whereas at the trough only 140 overlapped and the highest number of overlapping genes was shared between T2 vs. T0 and T4 vs. T0 (Fig. 3E). Among clock genes, *Clock*, *Cry2*, *Nr1d1*, *Nr1d2*, *Per1* and *Tef* were significantly affected by the heat pulse regardless of timing of treatment (Fig. 3F).

**Figure 3.**
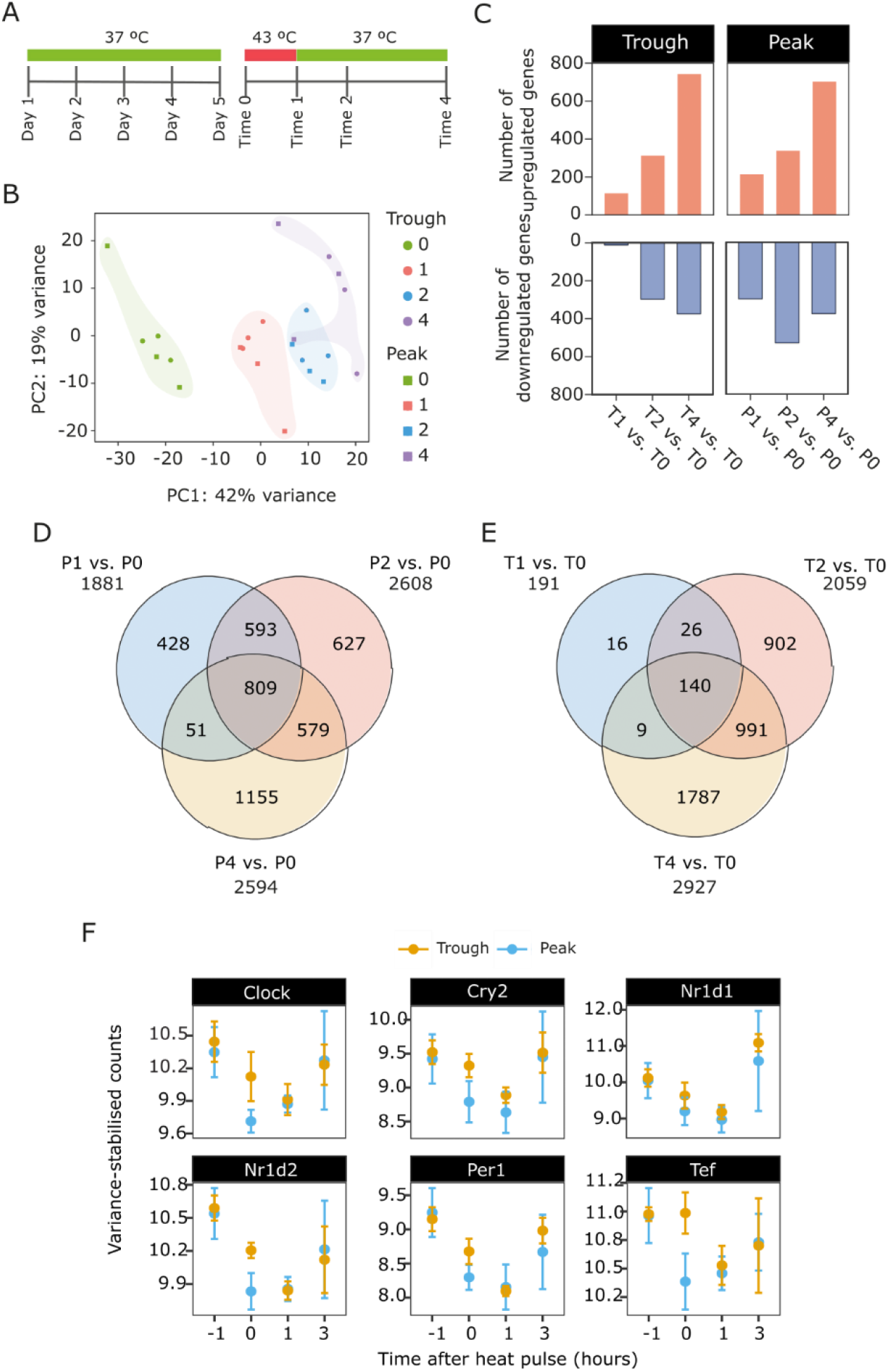
Heat pulse induces transcriptome wide response in mouse articular cartilage. **A.** RNA sequencing was performed on femoral head articular cartilage isolated from one-month-old mice. Explants were maintained at 37 °C for approximately five days. On the fifth day, samples that were snap frozen in liquid nitrogen before the heat pulse were labelled Time 0. Other samples were incubated at 43 °C for 60 min at a projected trough or peak of PER2::LUC, after which they were snap frozen at three time points: immediately after the heat pulse (Time 1), and one (Time 2) and three hours (Time 4) after the completion of the pulse. The numbers denote the number of hours that have passed since the first sample (Time 0) was collected. **B.** Principal component analysis of the RNA-seq samples **C.** Representation of the total number of up- and down-regulated in each pairwise comparison as bars (|log_2_ fold change| ≥1; FDR < 0.05). **D.** Venn diagram showing the overlap of heat-regulated genes (FDR < 0.05) amongst pairwise comparisons upon a heat pulse at a projected peak of PER2::LUC. **E**. Venn diagram showing the overlap heat-regulated genes (FDR < 0.05) amongst pairwise comparisons upon a heat pulse at a projected trough of PER2::LUC. In C, D and E the pairwise comparisons pertain to the trough and peak experimental conditions at different time points (Trough: T0, T1, T2 and T4; Peak: P0, P1, P2 and P4). FDR, false discovery rate. **F.** Expression levels of selected circadian clock genes in samples treated either at projected peak or trough of PER2::LUC.

Clustering of statistically significant genes from all comparisons coupled with gene ontology over-representation analysis provided insights into the functional relevance of different groups of heat-regulated genes (Supplementary Fig. 5 and Supplementary Table 1 and 2). Most clusters were associated with an up-regulation of gene expression and included biological process terms that related to the heat shock response, namely transcription by RNA polymerase II, RNA processing, ribonucleoprotein complex assembly and translation. Indeed, 45 genes encoding molecular chaperones were significantly modulated by the heat pulse (Fig. 4A). All the detected *Hsp* genes were significantly up-regulated and most of them reached their highest expression three hours after heat shock (Time 4). Moreover, 25 *Dnaj* family cochaperones were differentially regulated (Fig. 4B). The described transcriptional profile suggests that heat shock induces a cytoprotective response in chondrocytes, as expected.

**Figure 4.**
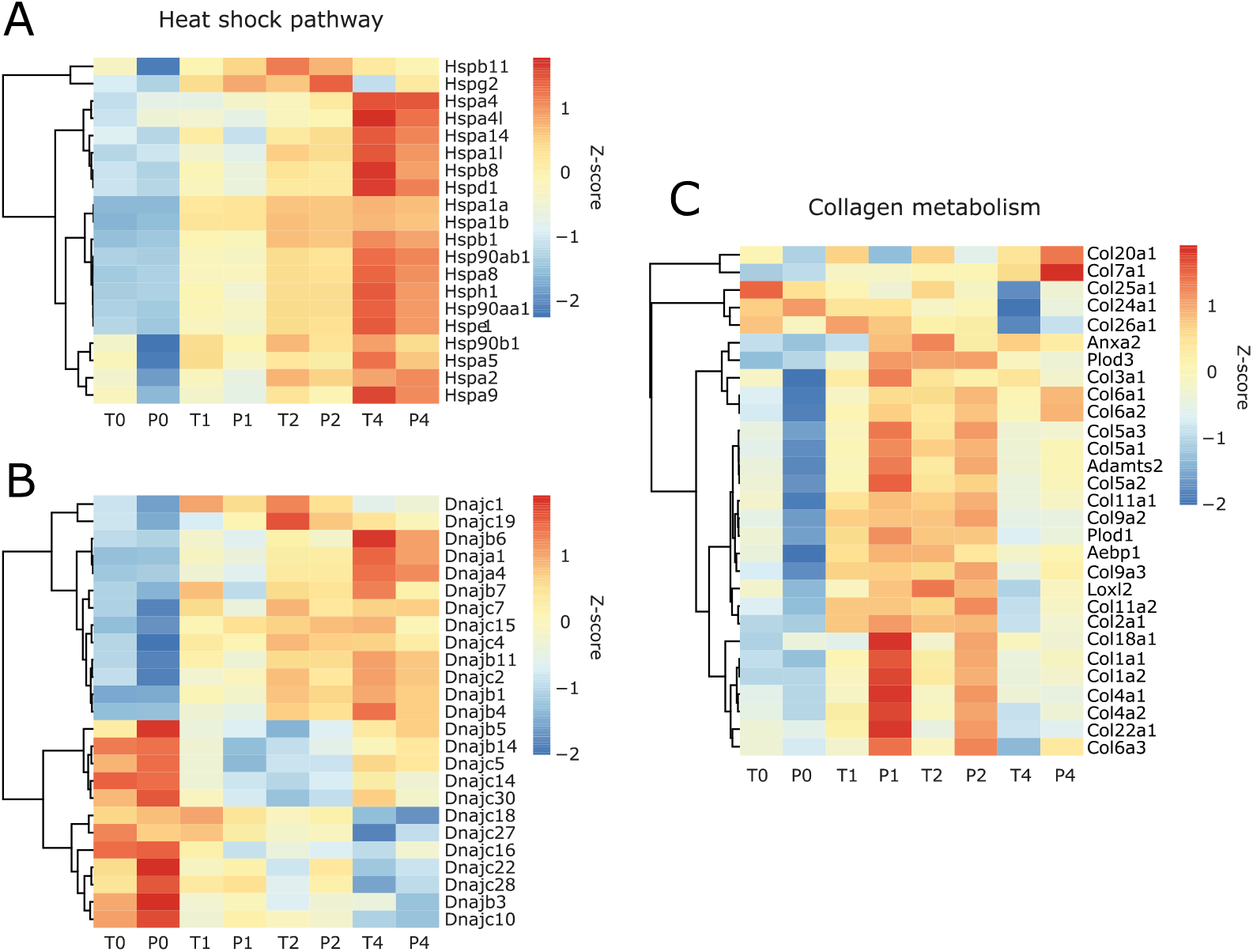
The heat pulse induced a cytoprotective and anabolic response in mouse articular cartilage. Heat map depicting response of heat shock proteins **A,** *Dnaj* family cochaperones **B**, and collagens **C** to the heat pulse. Trough: T0, T1, T2 and T4; Peak: P0, P1, P2 and P4. The mean of three biological replicates (n = 3) was presented. All presented genes are significantly differentially expressed (|log_2_ fold change| ≥1; FDR < 0.05).

Interestingly, several genes coding for proteins that are structural components of the extracellular matrix or partake in its assembly and remodelling were up-regulated upon heat shock (Fig. 4C). Of particular significance was the increase in the expression of the gene encoding collagen type II (*Col2a1)*. Genes that encode collagens III, VI, IX and XXII were also up-regulated as well as the anti-inflammatory *Anxa1*, matrix cross-linking *Loxl2* and *Adamts2*, a procollagen N-propeptidase (Fig. 4C) [39–41].

### Expression of heat-regulated and core clock genes in human osteoarthritic knee articular cartilage

To investigate whether heat pulses have comparable effect on human knee articular cartilage as on mouse tissues, qPCR was used to assess changes in the expression of most significantly up-regulated (*HSPB1*, *HSPA1A*, *HSPA1B*) and down-regulated (*JARID2*, *LNPEP*, *CXXC5*) genes that were identified in the mouse RNA-seq data. Cartilage explants dissected from the femoral condyles of three osteoarthritis patients were either maintained at 37 °C or heat shocked at 43 °C for 60 min and then returned to 37 °C. After the heat pulse, a strong induction in the expression of *HSPB1*, *HSPA1A* and *HSPA1B* was observed in samples from all donors in a pattern similar to the mouse RNA-seq data (Fig. 5A). Expression of *JARID2*, *LNPEP* and *CXXC5* was significantly down-regulated in at least one time point in at least 2 out of 3 heat-shocked samples (Fig. 5A). Expression of core clock genes *BMAL1*, *CLOCK*, *CRY1*, *CRY2*, *PER1* and *PER2* tended to show significant upregulation at least at one time point despite evident donor variability (Fig. 5B), indicating that similar heat pulse strategies could be effective in human cartilage to restore circadian clock rhythmicity.

**Figure 5.**
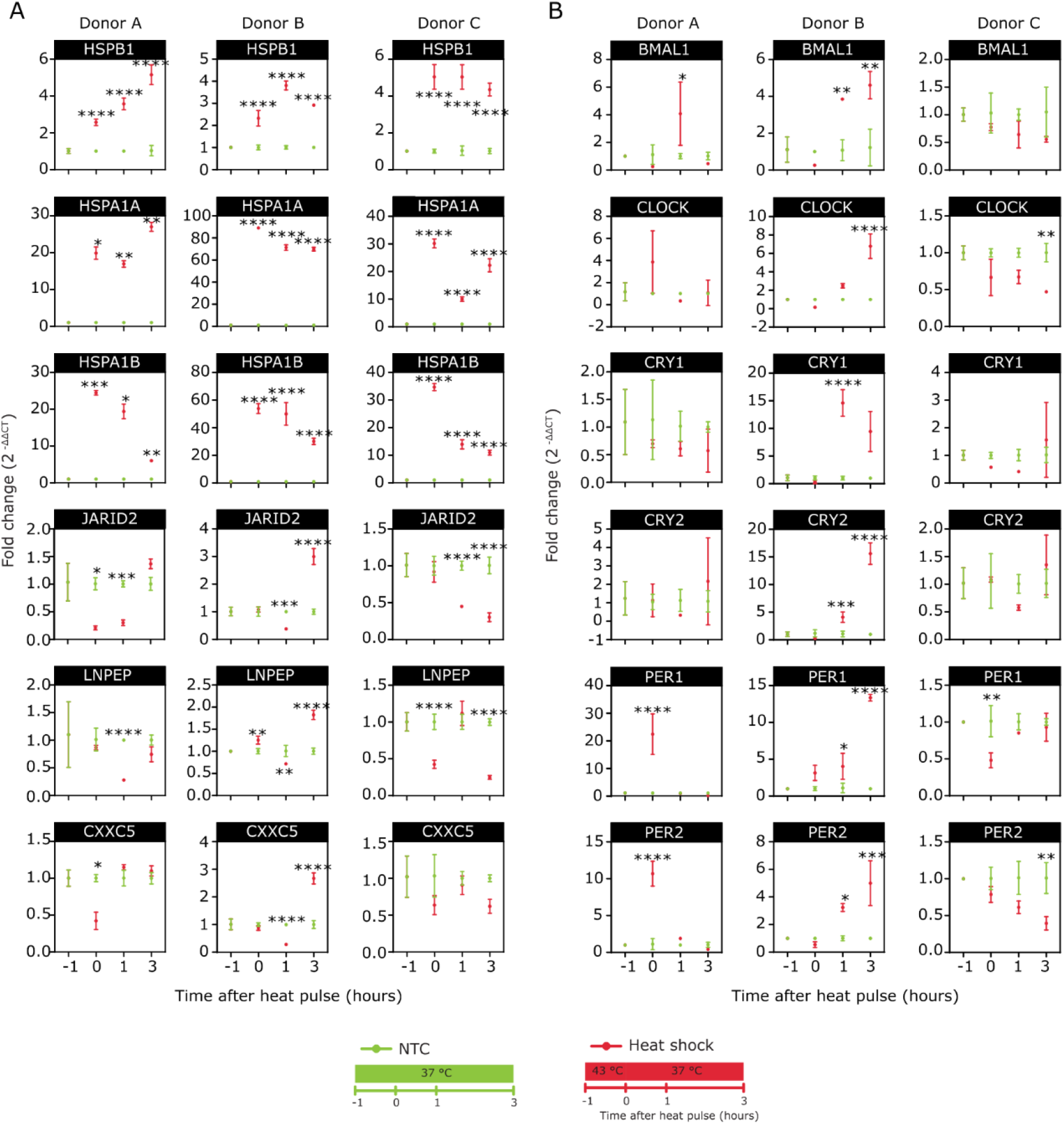
The heat pulse induced gene expression changes in human osteoarthritic knee articular cartilage. **A.** RT-qPCR quantification of *BMAL1*, *CLOCK*, *CRY1*, *CRY2*, *PER1* and *PER2* in articular cartilage isolated from human femoral condyles with osteoarthritis. Total RNA was extracted from cartilage explants incubated at 37 °C (NTC, no-treatment control) or heat shocked at 43 °C for 60 min and then returned to 37 °C (Heat shock). Data were first normalised to the mean expression of the reference genes CANX and CSDE1 and then to the NTC at time 0. **B.** RT-qPCR quantification of *HSPB1*, *HSPA1A*, H*PSA1B*, *JARDID2*, *LNPEP* and *CXXC5.* Data represent the mean ± SD (qPCR technical variation). Two-way ANOVA with a post-hoc Sidak’s multiple comparisons test (*, p ≤ 0.05; **, p ≤ 0.01; ***, p ≤ 0.001; ****, p ≤ 0.0001).

### Synchronisation of the circadian clock by a heat pulse is mediated by the HSP90 chaperone and the actin cytoskeleton

Previous studies have suggested that the heat shock response is involved in the regulation of circadian gene expression in peripheral tissues such as the liver and lungs [3,10,12]. This connection between the heat shock response and the circadian clock is further strengthened by the presence of HSEs (heat shock elements) in the promoter region of *Per2* [8]. Moreover, HSP90 has been reported to participate in circadian pacemaking in flies, plants and mammals [13,42,43]. To assess the role of the heat shock pathway on synchronisation of the clock, the effect of KNK437, a pan-HSP inhibitor that inhibits the synthesis of inducible HSPs (such as HSP105, HSP72 and HSP40) [44] and 17-DMAG, an analogue of geldanamycin that binds to the ATP binding site of HSP90 [45], were used. Pre-treatment with KNK437 had no effect on synchronisation of the chondrocyte clock by the heat pulse (Supplementary Fig. 6A-D). However, inhibition of HSP90 activity with 17-DMAG abolished the heat-induced resynchronization in a dose dependent manner (Fig. 6A and B), suggesting a significant role for HSP90 activity in the heat synchronisation process of chondrocytes.

**Figure 6.**
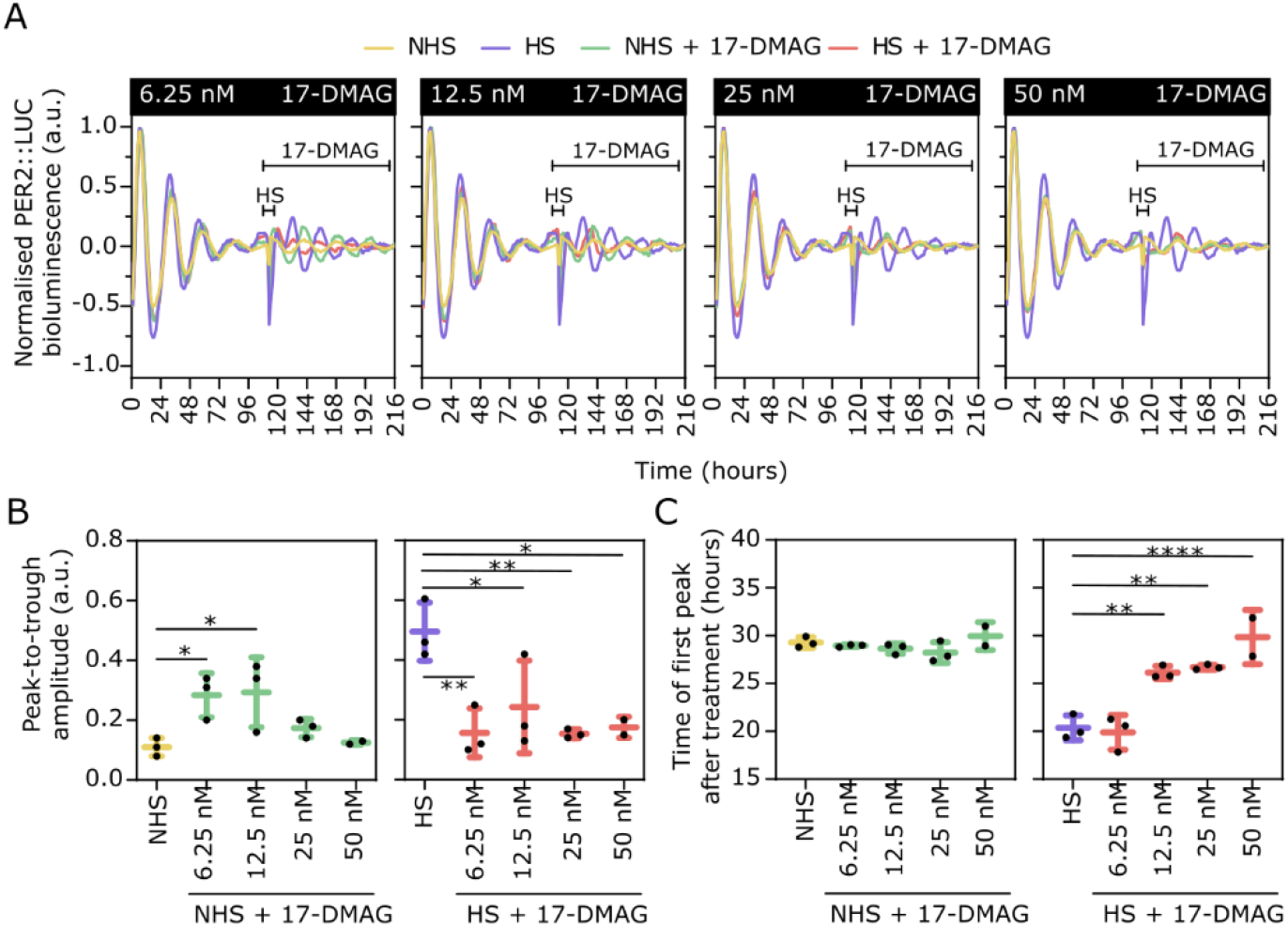
Inhibition of HSP90 activity with 17-DMAG blocked the heat induced resynchronisation of the circadian clock in PER2::LUC mouse primary chondrocytes. **A.** Normalised PER2::LUC signal from chondrocytes treated with 17-DMAG from 112 hours onwards. **B**. Peak-to-trough amplitude of the first oscillation after treatment. **C**. Time of the first peak following treatment. Data were reported as mean ± SD of three replicates. One-way ANOVA and Dunnett’s multiple comparisons test (*, p ≤ 0.05; **, p ≤ 0.01; ***, p ≤ 0.001; ****, p ≤ 0.0001). a.u – arbitrary units.

Actin is an essential component of the cytoskeleton that participates in the regulation of cell shape, movement and division, and cell signalling [16]. In mammalian peripheral clocks, actin plays a crucial role in regulation of the clock by mechanical properties of the extracellular matrix [17–19,46]. Additionally, heat stress can lead to the reorganisation and even collapse of actin networks [14,15]. To determine whether actin-mediated signalling is involved in the heat-induced resynchronisation of the circadian clock in chondrocytes, four different drugs affecting actin dynamics were tested: cytochalasin D (inhibitor of actin polymerisation), latrunculin B (inducer of actin depolymerisation), jasplakinolide (inducer of actin polymerisation) and Y-27632 (ROCK inhibitor). Pre-treatment of mouse primary chondrocytes with cytochalasin D was able to block the effect of the heat pulse on PER2::LUC amplitude, at all three tested concentrations, as well as phase shifting at higher concentration (Fig. 7A and B, Supplementary Fig. 7A). Pre-treatment with latrunculin B had a similar effect (Fig. 7C and D, Supplementary Fig. 7B). Interestingly, exposure to high concentration of jasplakinolide, which has opposite effect on actin compared to the previously used compounds, led to a significant amplitude increase in control cells mimicking the effect of the heat pulse (Fig. 7E and F, Supplementary Fig. 7C).

**Figure 7.**
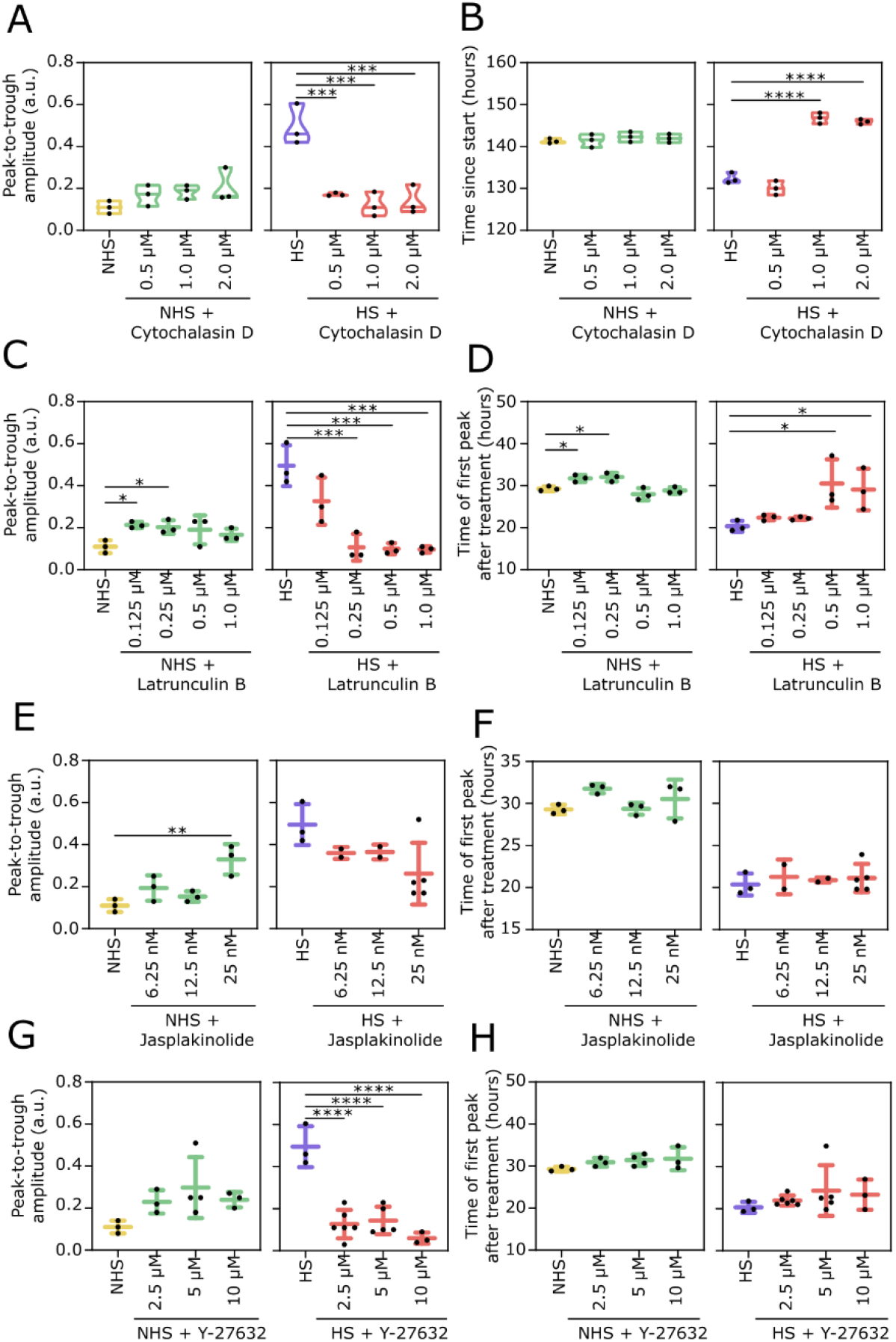
Inhibition of actin cytoskeleton reorganisation blocks heat induced resynchronisation of the circadian clock in PER2::LUC mouse primary chondrocytes. Quantification of normalised PER2::LUC signal from chondrocytes treated from 112 hours onwards with cytochalasin D (A, B), Latrunculin B (C, D), Jasplakonolide (E, F) and Y-27632 (G,H) in combination with heat pulse 60 min at 43 °C or NTC control. Peak-to-trough amplitude of the first oscillation after treatment (A, C, E, and G). Time of the first peak following treatment (B, D, F and H). Data reported as mean ± SD of three replicates. One-way ANOVA and Dunnett’s multiple comparisons test (*, p ≤ 0.05; **, p ≤ 0.01; ***, p ≤ 0.001; ****, p ≤ 0.0001). a.u – arbitrary units.

ROCK proteins are critical Rho GTPases that regulate the assembly, disassembly and dynamic rearrangements of the actin cytoskeleton [47,48]. In non-heat-shocked chondrocytes, chronic incubation with Y-27632 did not significantly alter the amplitude of the first oscillation or the timing of the first peak. However, the ROCK inhibitor completely blocked the synchronising effect of heat pulse, suggesting the preservation of filamentous actin structures is important to resynchronise the clock following heat pulse (Fig. 7G and H, Supplementary Fig. 7D).

## Discussion

Circadian clocks generate daily rhythms in molecular, cellular, physiological and behavioural functions, providing a temporal dimension to organismal homeostasis. With ageing, the musculoskeletal system shows profound functional decline, including loss of bone and degeneration of cartilage, leading to frailty and loss of mobility [49,50]. The circadian clock, with hundreds of genes under its rhythmic control, is important in cartilage homeostasis [6]. Circadian oscillations in gene expression dampen in ageing cartilage [4,34], therefore new approaches are needed to restore circadian pacemaking and slow down tissue ageing. Here, we systematically investigated the effect of heat pulse as a new way of re-synchronising cartilage clocks. Our data show that one hour heat pulse at 43°C at the peak of daily PER2 expression strongly induces circadian clock oscillations in cartilage explants and cultured chondrocytes. Mechanistically, HSP90 and the actin-Rho/ROCK pathways are involved in the heat-induced resynchronisation of the clock.

Recent studies propose a bidirectional relationship between circadian clocks and ageing. Many of the hallmarks of ageing, such as deregulated nutrient sensing, mitochondrial dysfunction, slowed cell division rates, loss of proteostasis, altered intercellular communication and disrupted mechanosensing of ECM stiffness, either affect circadian clock function and/or are governed by the clock [46]. Notably, improvement of circadian rhythms may delay age-related functional decline, opening novel and exciting research avenues into clock-based therapeutic strategies for degenerative diseases such as hypercholesterolemia, hypertension, endocrine disorders, cancers and rheumatic diseases [51–53]. Our data showed that in both cells and explants (from young and ageing mice), a 60- min pulse at 43 °C was sufficient to restore dampened circadian rhythms. Moreover, a time-of-day-dependent PER2::LUC response to heat shock was detected in mouse cartilage explants. Specifically, heat pulse elicited the strongest response at the peak of PER2 expression which in mouse cartilage *in vivo* coincides with the onset of activity. Interestingly, we have recently shown that mechanical loading resets the clock in cartilage with the strongest clock amplitude enhancing effect at the same phase of the clock as the heat pulse [34]. During loading of viscoelastic tissues, part of the mechanical energy is transformed into heat that can locally increase the tissue temperature. This is the case in cartilage where temperature can increase from resting 31°C to 37°C after 60 min of jogging [54]. Combination of mechanical loading and temperature was recently shown to have synergistic effect on chondrogenesis *in vitro* [55], suggesting a strong connection between cartilage biomechanical properties and daily changes in physiology and homeostasis. Importantly, the heat pulse was able to resynchronise the circadian clock in aged mouse cartilage explants, and in cartilage explants with the presence of the clock disrupting cytokine IL-1α. Preservation of the circadian clock oscillations was recently shown to be beneficial in preventing loss of extracellular matrix after treatment with proinflammatory cytokines in engineered cartilage constructs [38].

RNA sequencing offered a crucial view of the transcriptional reprogramming that occurs in mouse articular cartilage in the first three hours following exposure to high temperature. Specifically, a 60- min pulse at 43 °C elicited a cytoprotective response that was characterised by the extensive up-regulation of processes related to transcription, translation as well as heat shock factors and chaperones. This is in line with literature where although global translation is generally suppressed in response to most cellular stresses, subsets of transcripts that produce proteins vital for cell survival and stress recovery are selectively translated [56]. One of the best-known examples of this translational control involves the HSP family, whose mRNA synthesis is up-regulated in cells upon exposure to elevated temperature [57]. The cytoprotective response mounted by HSPs allows cells, tissues or intact organisms to survive exposure to elevated temperatures, conferring protection against subsequent and otherwise lethal hyperthermia [14].

In mammals, it has been demonstrated that HSF1 is essential for the synchronisation of *Per2*/PER2 rhythms in fibroblasts by a 30-min pulse at 43 °C and a physical association between HSF1 and BMAL1 was detected [8]. Through pharmacological experiments with KNK437 (inhibitor of several HSPs, such as HSP105, HSP70, and HSP40), Buhr *et al.* demonstrated that the heat shock pathway is integral to temperature resetting and inhibition of these HSPs eliminated phase shifting of the clock in lung and pituitary in response to temperature changes [3]. In our experiments, pre-treatment of chondrocytes with 17-DMAG (inhibitor of HSP90 activity) abolished the synchronisation of the clock in response to the heat pulse. However, unlike in Buhr *et al.* [3], the same was not observed with KNK437 in all tested concentrations, suggesting some of these mechanisms might be cell- and/or tissue-specific.

Signalling through the actin cytoskeleton and SRF pathway has been shown to be involved in the clock resetting effects of serum shock [58] as well as the effect of extracellular matrix stiffness on the clock [19], while heat shock has been observed to disrupt the actin cytoskeleton [15]. Connecting these two lines of evidence, here we demonstrated that drugs that destabilise filamentous actin significantly blocked the heat-mediated resynchronisation of the clock, while jasplakinolide, which induces actin polymerisation, did not block the heat-mediated resynchronisation of the clock. Moreover, inhibition of the actin associated Rho/ROCK signalling pathway prevented resynchronisation of the chondrocyte clock. This evidence establishes a direct connection between the heat-mediated perturbation of an important cellular structure, actin cytoskeleton, and the resynchronisation of the chondrocyte clock under conditions of physiological stress.

Given that a significant portion of cartilage physiology is under circadian control, and chronic disruption of the clock rhythm can lead to cartilage tissue degeneration, our findings of heat resynchronisation of the cartilage clock could be utilised as a potential therapy to slow down cartilage degradation in the context of osteoarthritis. Of note, thermotherapy is frequently used in clinical settings as an adjunctive therapy to alleviate pain in osteoarthritic joints [59,60]. However, the vast majority of studies on the benefits of thermotherapeutic treatment in musculoskeletal disorders (e.g., osteoarthritis and rheumatoid arthritis) lack standardisation and an adequate number of patients [61,62], with little appreciation of treatment timing. Although our study was limited to only three osteoarthritic cartilage donors, it is encouraging that the experiments show induction of heat shock response and induction of circadian clock genes in human osteoarthritic cartilage explants. Nevertheless, further investigation and clinical trials are needed to establish the usefulness of time-prescribed heat pulses as an intervention in a clinical setting.

## Funding

Versus Arthritis Senior Research Fellowship Award 20875 (QJM); Wellcome Trust MCB PhD Studentship 215205/Z/19/Z (CFG); MRC project grants MR/T016744/1 and MR/P010709/1 (QJM, JAH); MRC Confidence in Concept grant (QJM, LB); Wellcome Trust ‘Projects for Translation’ Award (QJM, LB); BBSRC sLoLa grant BB/T001984/1 (QJM).

## Acknowledgments

We thank J Takahashi (UT Southwestern Medical Center, US) for the PER2::LUC mouse line. We thank the Genomics Core Facility (A Hayes and L Zeef) for their kind assistance with RNAseq. We also thank the Bioimaging facility and David Spiller for assistance with microscopy.

## Supplementary Figures

**Supplementary Figure 1.**
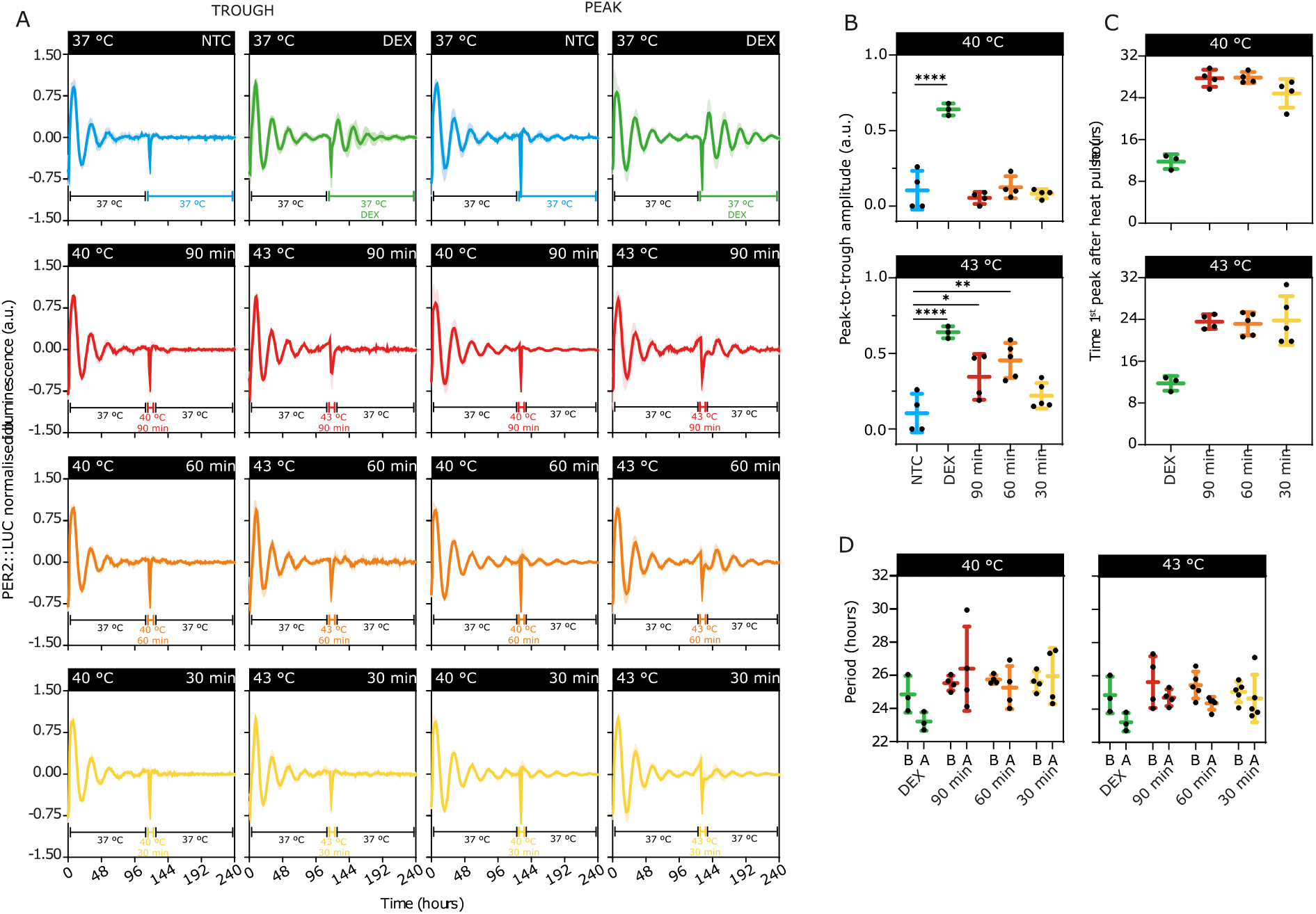
Dampened circadian rhythms in murine articular cartilage can be restored by heat pulses. **A.** Normalised bioluminescence profiles of mouse femoral head articular cartilage. Explants were treated either at a projected trough (left) or peak (right) of PER2::LUC. Bioluminescence profiles showed an increase in amplitude when tissues were treated at a projected peak, whilst temperature pulses at a projected trough did not restore circadian oscillations. **B.** Peak-to-trough amplitude of the first oscillation after treatment compared to the no-treatment control (NTC). Data were normalised to the maximum bioluminescence value for each individual trace. 40 °C: one-way ANOVA (*p* < 0.0001) and post-hoc Dunnett’s multiple comparison test (****, *p* < 0.0001); 43 °C: one-way ANOVA (*p* < 0.0001) and post-hoc Dunnet’s multiple comparison test (****, *p* < 0.0001; **, *p* = 0.0010; *, *p* = 0.0276). **C.** Time of the first peak (circadian phase) following treatment. **D.** Period of the circadian rhythms before (B) and after (A) treatment. Data were reported as mean ± SD. n ≥ 3 in all experiments. DEX, 100 nM dexamethasone; a.u., arbitrary units.

**Supplementary Figure 2.**
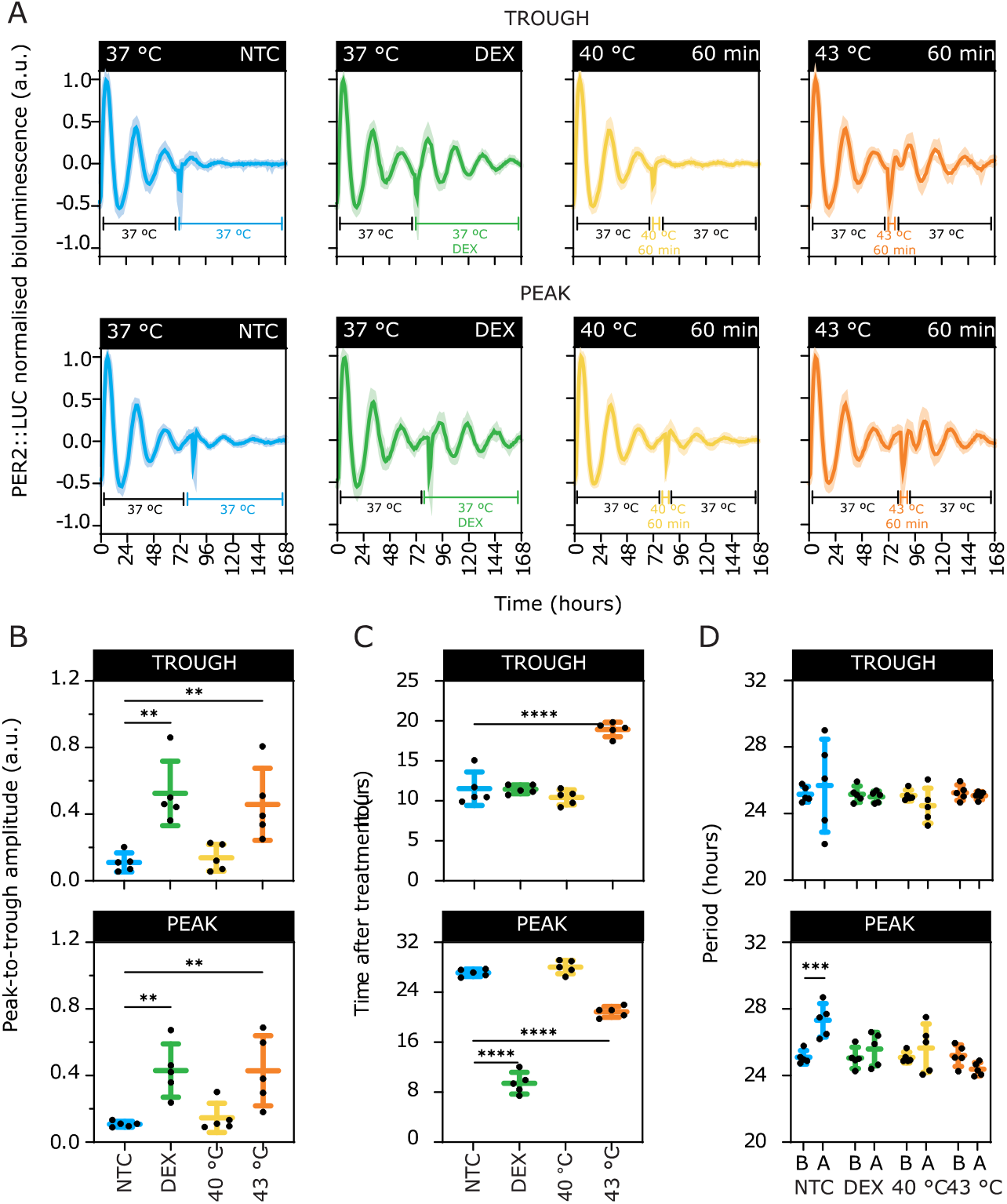
Effect of a heat pulse at a PER2::LUC trough (72 hours) or peak (84 hours) on the circadian rhythms of mouse articular chondrocytes. **A.** Normalised bioluminescence profiles of primary chondrocytes. Cells were treated either at a trough (72 hours) or peak (84 hours) of PER2::LUC. **B.** Peak-to-trough amplitude of the first oscillation after treatment compared to the no-treatment control (NTC). One-way ANOVA and post-hoc Dunnet’s multiple comparison test. Trough, *p* = 0.0007; DEX: *p* = 0.0016; 43 °C: *p* = 0.0066. Peak: *p* = 0.0019; DEX, *p* = 0.0058; 43 °C, *p* = 0.0060. **C.** Time of the first peak following treatment compared to the NTC. One-way ANOVA and post-hoc Dunnett’s multiple comparison test. Trough: *p* < 0.0001; ****, *p* < 0.0001. Peak: *p* < 0.0001; ****, *p* < 0.0001. **D.** Period of the circadian rhythms before (B) and after (A) treatment. Two-way ANOVA and Šídák’s multiple comparisons test. *p* = 0.0050; ***, *p* < 0.0001. Data were reported as mean ± SD. n = 5 in all experiments. DEX, 100 nM dexamethasone; a.u., arbitrary units.

**Supplementary Figure 3.**
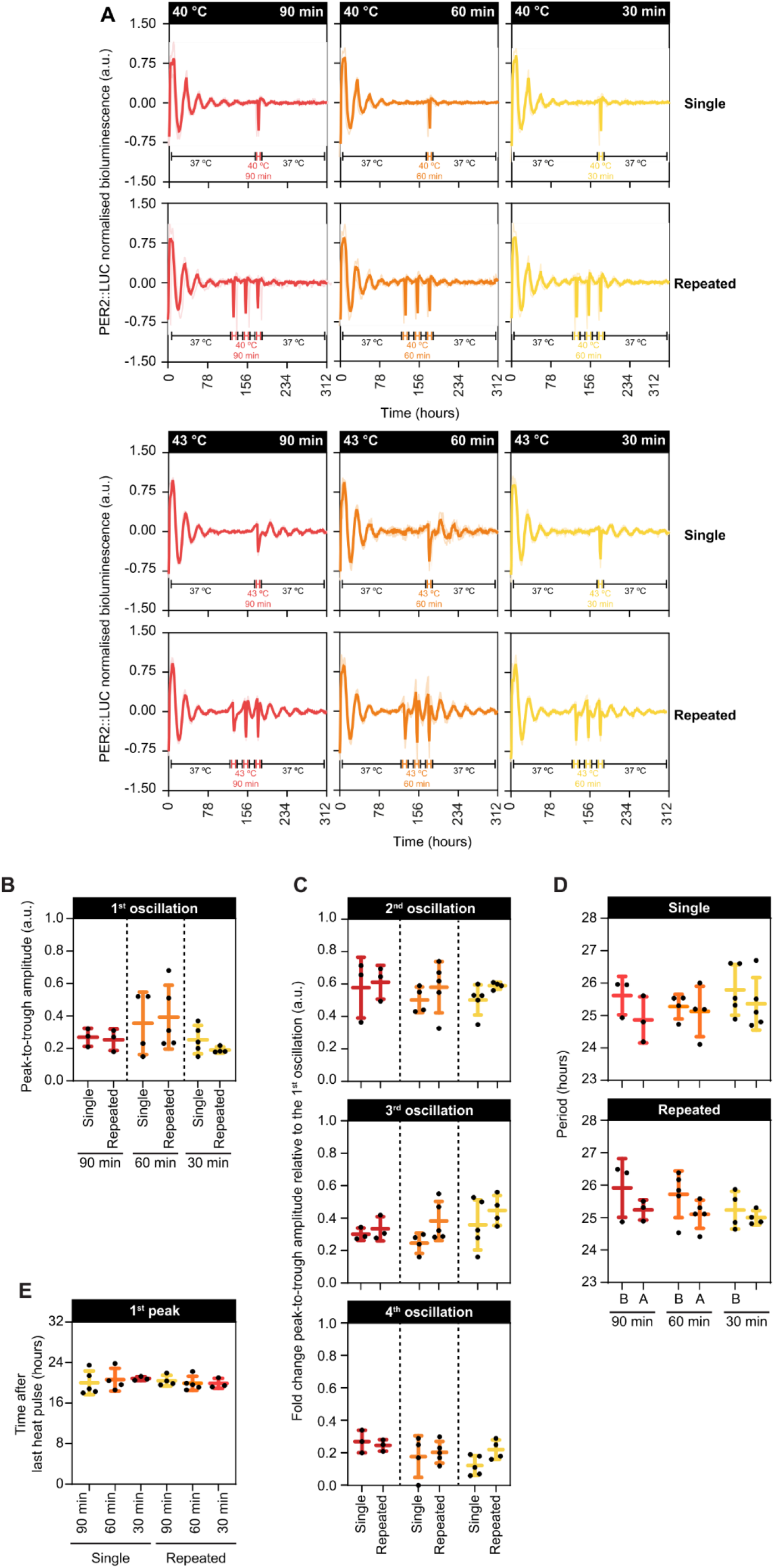
Repeated heat pulses at a projected peak of PER2::LUC do not further improve the amplitude of circadian rhythms in mouse articular cartilage. **A.** Normalised bioluminescence profiles of femoral head articular cartilage from PER2::LUC reporter mice. Heat shocks at 40 °C (upper panel) or 43 °C (lower panel) were applied once on the seventh day (top) or repeated on three consecutive days (bottom) starting on the fifth day of *ex vivo* culture. Explants were heat shocked at a projected peak of PER2::LUC with increasing duration (30-90 min). **B.** Peak-to-trough amplitude of the first oscillation after the last heat pulse. **C.** Fold change in the peak-to-trough amplitude normalised to the first oscillation of the respective experimental condition. **D.** Period of the circadian rhythms before (B) and after (A) treatment. **E.** Time of the first peak following heat treatment. Data were reported as mean ± SD. n ≥ 3 in all experiments.

**Supplementary Figure 4.**
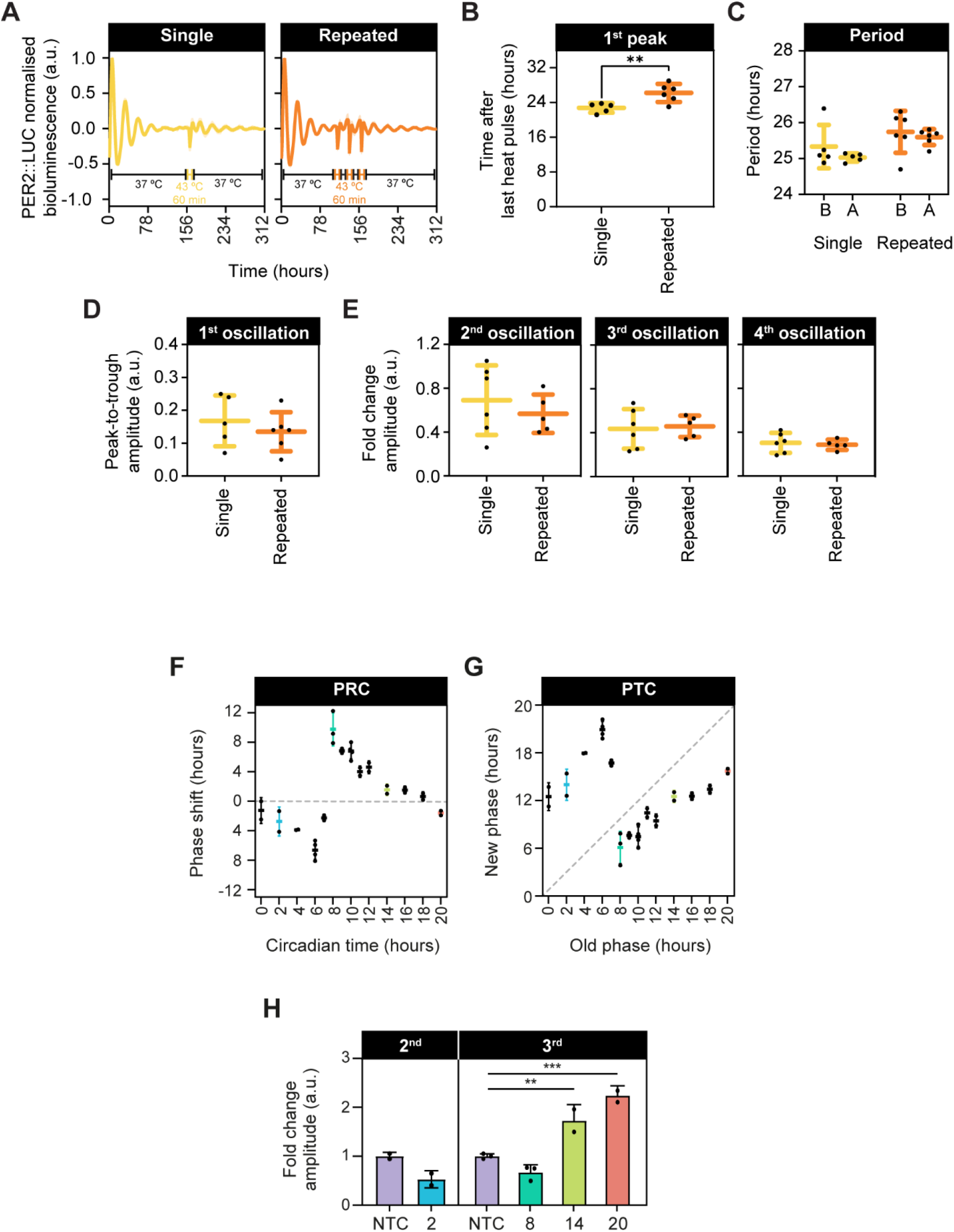
Repeated heat pulses at a projected peak of PER2::LUC do not further improve the amplitude of circadian rhythms in mouse primary chondrocytes. **A**. Normalised bioluminescence profiles of mouse primary chondrocytes from PER2::LUC reporter mice. Heat shocks at 43 °C were applied once on the sixth day or repeated on three consecutive days starting on the fourth day of culture. **B**. Time of the first peak following heat treatment. Two-tailed unpaired t-test, *p* = 0.0092). **C**. Period of the circadian rhythms before (B) and after (A) treatment. **D.** Peak-to-trough amplitude of the first oscillation after the last heat pulse. **E**. Fold change in the peak-to-trough amplitude normalised to the first oscillation of the respective experimental condition. **F.** Phase response curve for mouse primary chondrocytes in response to a 60-min pulse at 43 °C. Phase shifts were determined by subtracting the time of the peak after treatment to the time of observed peaks in the no-treatment control (NTC). **G.** Phase transition curve showing the phase of circadian rhythms in chondrocytes before (old phase) and after (new phase) the heat pulse. The dashed line segment represents a slope equal to one. **H.** Peak-to-trough amplitude of the first oscillation after heat shock compared to the respective NTC oscillation as indicated. Second oscillation: two-tailed unpaired t-test (*p* = 0.0744). Third oscillation: one-way ANOVA (*p* = 0.0003) and post-hoc Dunnett’s multiple comparison test (NTC vs. CT14, *p* = 0.0115; NTC vs. CT20, *p* = 0.0007). Data were reported as mean ± SD. n ≥ 5 in A-E and n ≥ 2 in F-H.

**Supplementary Figure 5.**
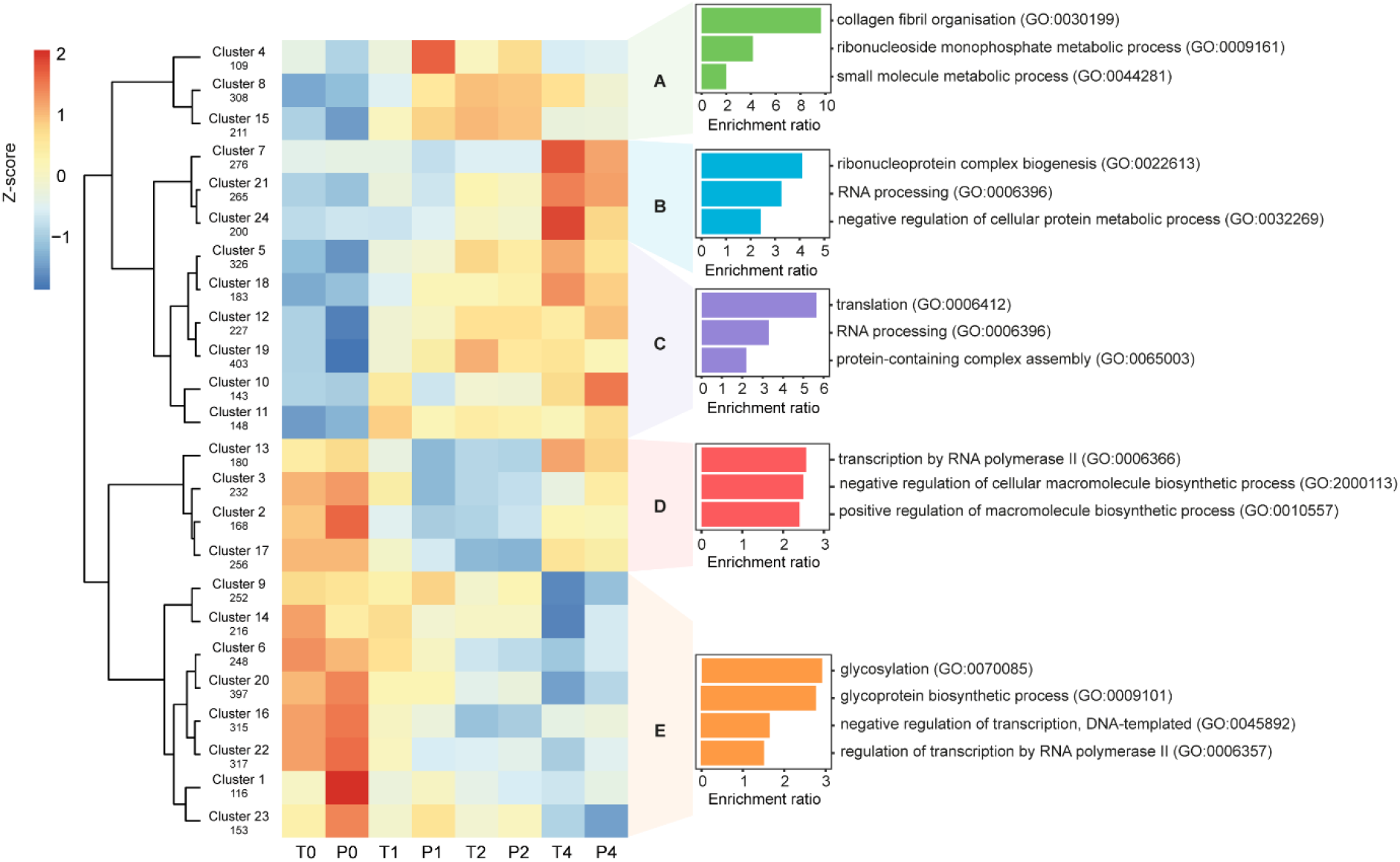
Heatmap of k-means clustering of statistically significant genes and gene ontology over-representation in mouse primary chondrocytes. The batch-corrected read counts of the 5,650 statistically significant genes (FDR < 0.05) were standardised to build a heatmap (Z-score). Pearson correlation was used as the dissimilarity metric; and the colours in the map represent row-scaled expression values: blue, yellow and red are associated with low, intermediate and high expression levels, respectively. Cluster size is indicated underneath its name. A gene over-representation analysis was performed to investigate the biological function of selected groups of clusters. The most over-represented gene ontology (GO) terms pertaining to biological process were shown as well as their enrichment ratio. Trough: T0, T1, T2 and T4; Peak: P0, P1, P2 and P4). The mean of three biological replicates was presented for each time point (n = 3). FDR, false discovery rate.

**Supplementary Figure 6.**
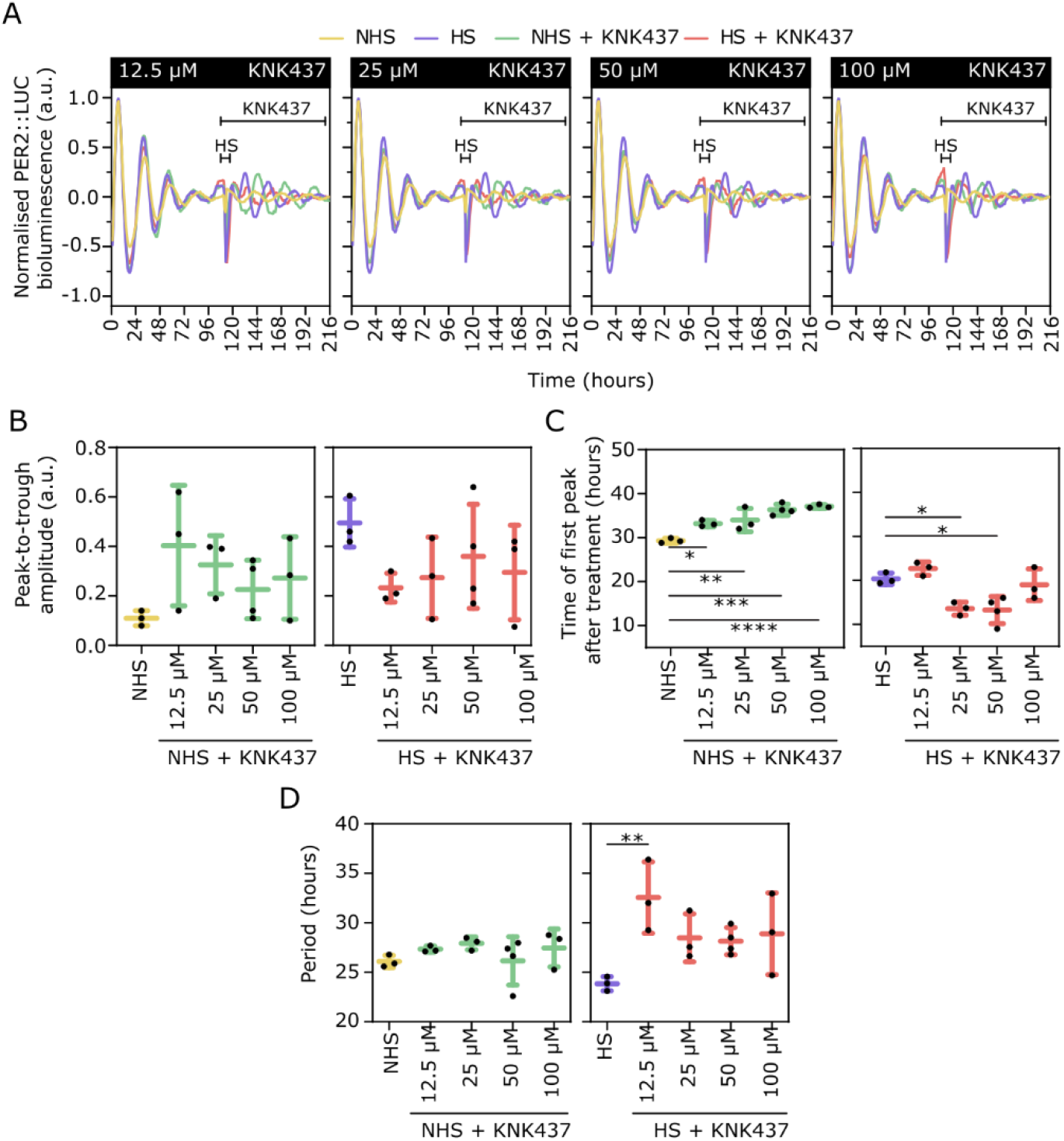
The pan-HSP inhibitor KNK437 did not block the heat pulse induced synchronisation of the circadian clock in mouse primary chondrocytes. **A**. Normalised PER2::LUC signal from chondrocytes treated with KNK437 from 112 hours onwards. **B**. Peak-to-trough amplitude of the first oscillation after treatment. **C**. Time of the first peak following treatment. One-way ANOVA and Dunnett’s multiple comparisons test. NHS: *p* = 0.0002; *, *p* = 0.0163; **, *p* = 0.0049; ***, *p* = 0.0001; ****, *p* < 0.0001. HS: *p* = 0.0015; 25 μM: *, *p* = 0.0204; 50 μM, *, *p* = 0.0103. **D**. Circadian period after treatment. One-way ANOVA and Dunnett’s multiple comparisons test. *p* = 0.03; **, *p* = 0.0070. Data in **B**, **C** and **D** were compared to the respective no-KNK437 control. n ≥ 3. a.u., arbitrary units. Data were reported as mean ± SD.

**Supplementary Figure 7.**
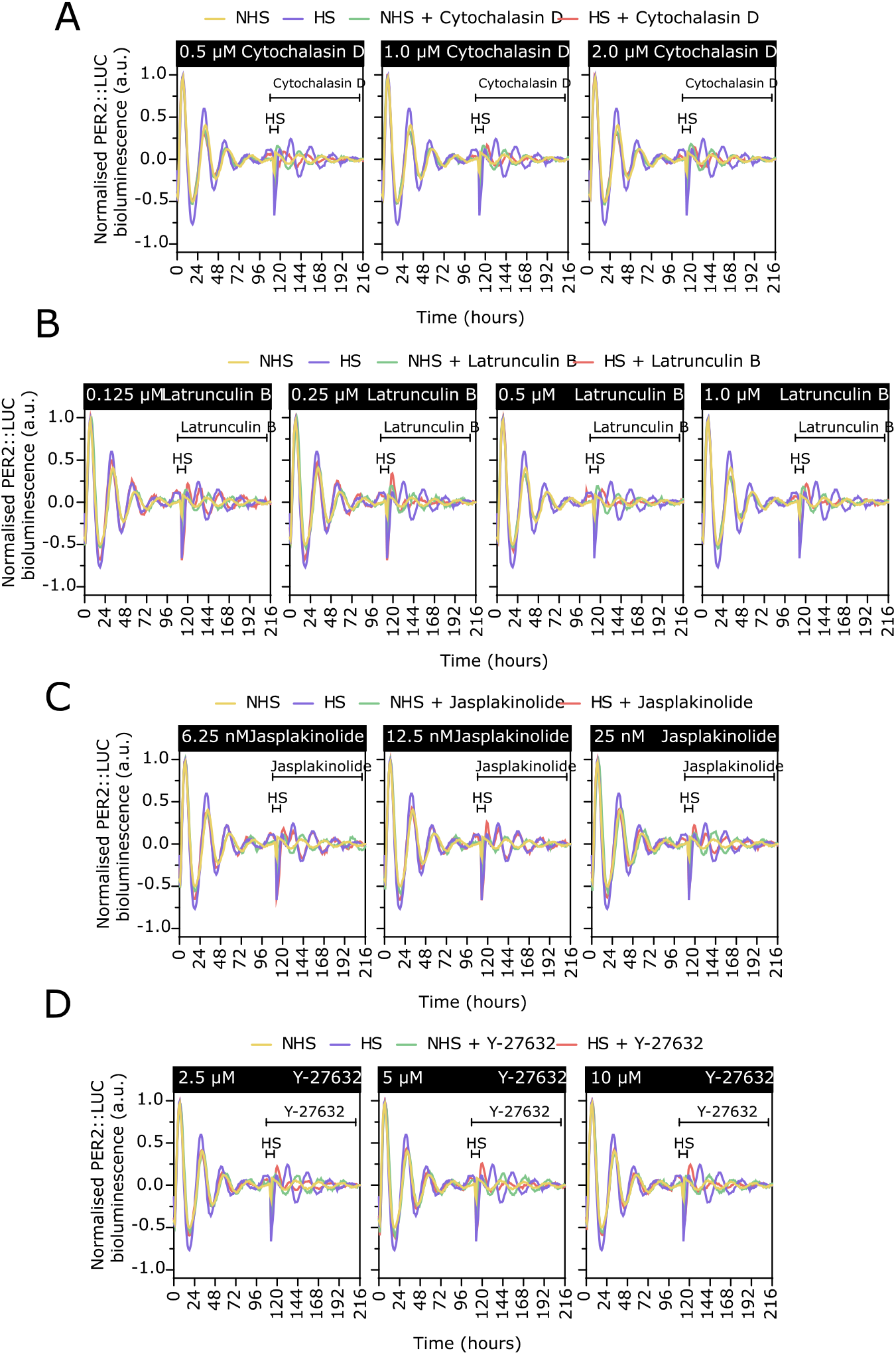
Inhibition of actin cytoskeleton reorganisation blocked the heat induced resynchronisation of the circadian clock in PER2::LUC mouse primary chondrocytes. Normalised PER2::LUC signal from chondrocytes treated from 112 hours onwards with Cytochalasin D **(A)**, Latrunculin B **(B)**, Jasplakonolide **(C)** and Y-27632 **(D)** in combination with a heat pulse of 60 min at 43 °C or NTC control.

## Notes

### Competing Interest Statement

The authors have declared no competing interest.

